# Dynamic contrast enhanced MRI with clinical hepatospecific MRI contrast agents in pigs: initial experience

**DOI:** 10.1101/2020.02.18.946541

**Authors:** Jeremy M.L. Hix, Christiane L. Mallett, Matthew Latourette, Kirk A. Munoz, Erik M. Shapiro

## Abstract

Pigs are an important translational research model for biomedical imaging studies, and especially for modeling diseases of the liver. Dynamic contrast enhanced (DCE)-MRI is experimentally used to measure liver function in humans, but has never been characterized in pig liver. Here we performed DCE-MRI of pig liver following the delivery of two FDA approved hepato-specific MRI contrast agents, Gd-EOB-DTPA (Eovist) and Gd-BOPTA (Multihance), and the non-hepatospecific agent Magnevist, and optimized the anesthesia and animal handling protocol to acquire robust data. A single pig underwent 5 scanning sessions over six weeks, each time injected at clinical dosing either with Eovist (twice), Multihance (twice) or Magnevist (once). DCE-MRI was performed at 1.5T for 60 minutes. DCE-MRI showed rapid hepatic MRI signal enhancement following IV injection of Eovist or Multihance. Efflux of contrast agent from liver exhibited kinetics similar to that in humans, except for one hyperthermic animal where efflux was very fast. As expected, Magnevist was non-enhancing in the liver. The hepatic signal enhancement from Eovist matched that seen in humans and primates, while the hepatic signal enhancement from Multihance was different, similar to rodents and dogs, likely the result of differential hepatic organic anion transport polypeptides. This first experience with these agents in pigs provides valuable information on contrast agent dynamics in normal pig liver. Given the disparity in contrast agent uptake kinetics with humans for Multihance, Eovist should be used in porcine models for biomedical imaging. Proper animal health maintenance, especially temperature, seems essential for accurate and reproducible results.

## Introduction

Pigs are a valuable translational biomedical tool, particularly for modeling human liver disease. Robust porcine models of cirrhosis ^1-4^, non-alcoholic steatohepatitis ^5, 6^, hepatocellular carcinoma ^7, 8^, diabetes ^9-11^, hereditary tyrosinemia type 1 ^12-15^ and acute liver failure ^16, 17^ have all been reported and mimic many clinical manifestations of the human disease. Biomedical imaging can be used in conjunction with these porcine models in many ways. For example, because pigs have similar anatomy (size, organization and location of organs) and physiology to humans, new biomedical imaging protocols can be developed in pig models of disease to better diagnose humans with disease, both at an earlier stage of disease and to follow treatment. Further, biomedical imaging can provide imaging biomarkers to support research and development for treatments for these diseases.

There are two FDA-approved MRI contrast agents that exhibit uptake in the human liver, Gd-EOB-DTPA (Eovist) and Gd-BOPTA (Multihance), mediated by organic anion transport polypeptides (OATP) OATP1B1 and OATP1B3 ^18^. In humans, Eovist exhibits rapid hepatic uptake with ∼ 50:50 clearance by the healthy liver and kidney following IV injection ^18^, while Multihance exhibits low hepatic uptake with ∼ 5:95 clearance by the healthy liver and kidney ^19^. Despite the different pharmacokinetics, both hepatospecific MRI contrast agents enable useful clinical MRI examinations, not only discriminating anatomic disease features in the liver, such as tumors, but also enabling the measurement of hepatic functional deficits.

While extensive data on hepatic uptake of these agents exists in humans and rodents (uptake transporters OATP1A1 and OATP1B2) ^20, 21^, and to some extent in dogs and rabbits (uptake transporter OATP1B4) ^22-24^, there is no data on the hepatic uptake of these two hepatospecific MRI contrast agents in pigs (uptake transporter OATP1B4). Further, given the high sensitivity of animal physiology for hepatic function, proper animal handling protocols for pigs need to be established ^25^. Given the aforementioned importance of pigs for modeling human liver disease and the potential for biomedical imaging in this regard, the narrow goals of this work were to 1) perform dynamic contrast enhanced (DCE)-MRI studies in pigs using these two hepatospecific MRI contrast agents, providing baseline assessments for contrast agent hepatic influx in normal pig liver, and 2) to determine optimal experimental animal protocols for acquiring high quality DCE-MRI data.

## Experimental

### General experimental paradigm

All animal experiments were performed under Michigan State University IACUC approved procedures. One pig underwent 5 separate 60 minute duration DCE-MRI sessions on the Michigan State University College of Veterinary Medicine (CVM) Siemens Esprit 1.5T MRI over a course of 6 weeks, each time injected with a different MRI contrast agent, according to Table 1. Equivalent human clinical gadolinium dose was administered each session for each contrast agent, which is different depending on the agent. Experiments were staggered by 1 or 2 weeks to ensure complete elimination of the agent in between sessions and for scheduling considerations.

**Table 1:** Contrast agent information and dosing for MRI studies.

| Scan session | Contrast agent & trial | Gadolinium concentration (mM) | Animal weight (kg) | Volume injected (ml) | Total gadolinium dose (micromoles) |
| --- | --- | --- | --- | --- | --- |
| 1 – Week 1 | Eovist 1 | 0.25 | 26.0 | 2.6 | 0.65 |
| 2 – Week 2 | Multihance 1 | 0.5 | 29.0 | 4.8 | 2.4 |
| 3 – Week 3 | Magnevist | 0.5 | 35.3 | 7.1 | 3.5 |
| 4 – Week 4 | Multihance 2 | 0.5 | 41.3 | 8.3 | 4.2 |
| 5 – Week 6 | Eovist 2 | 0.25 | 56.5 | 5.7 | 1.4 |

### Animal care

A 2.5 month old, 25 kg Yorkshire/PIC327 Crossbred male pig was purchased from Michigan State University Swine Farm and single-housed in standard large animal research conditions. No adverse events, injuries or infections were encountered during the course of animal housing.

### Anesthesia and animal maintenance during MRI

In the first scan session, the pig was lightly sedated via intramuscular (IM) Telazol (4 mg/kg) in the vivarium and carted to the MRI suite for intubation and anesthesia. This presented challenges for intubation so an easier regimen was designed for scan sessions 2-5 where the animal was sedated, anesthetized and intubated in the vivarium, and then wheeled under inhaled anesthesia to the MRI. Bilateral ear vein catheters were also installed in the vivarium and blood was drawn pre- and post-MRI to measure clinical chemistry. Intravenous (IV) lidocaine as a single injection (1 mg/kg) and IV Propofol as needed (2 mg/kg) were used to aid with intubation and animal transport.

During MRI, anesthesia was maintained via 2-3% sevoflurane in 100% oxygen with the pig breathing spontaneously rather than ventilated. The animal was placed inside the MRI bore in a chest-down, ventral recumbency. IV Fluid Therapy (0.9% NaCl for Injection) was given as a continuous drip (5-10 mL/kg/hr) during the MRI. Vital signs were stabilized and monitored during the MRI anesthetic event, which included echocardiogram, heart rate, respiratory rate, end-tidal CO_2_, blood pressure, and oxygen hemoglobin saturation. Core body temperature (CBT) was measured via rectal thermometer pre- and post-MRI. During scan session 2, the pig exhibited hyperthermia at the end of the experiment, resultant from the experiment and not malignant hyperthermia; therefore, in subsequent scan sessions 3-5, a cooling recirculating water pad was placed ventrally, under the animal. To further enable cooling, a plastic cover for the scanner bed used for scan sessions 1 and 2, was not used in scan sessions 3-5. These two mitigations maintained the animal at stable, normothermic CBT in scan sessions 3-5.

### MRI protocol

Following scout images to define anatomy, DCE-MRI was performed using a T_1_-weighted multi-slice gradient echo VIBE sequence with Cartesian k-space filling. Scan parameters were: TR = 3.65 ms, TE = 1.77 ms, flip angle = 12°, matrix size = 256 x 256, FOV = 32 x 32 cm, slice thickness = 3 mm, number of slices = 32, temporal resolution = 1 frame every 13.6 seconds, GRAPPA acceleration = 2.

After the fifth TR, rapid injection (delivered in < 2 s) of contrast agent according to Table 1 was performed as a manual IV bolus via ear vein catheter, followed by ∼10 mL manual IV bolus saline flush. Following DCE-MRI, an anatomical image of the abdomen was acquired to confirm accumulation of the agent into the bladder. Following all but the last MRI, the pig was recovered, extubated, and returned to the vivarium. Following the last MRI, while under deep inhalant anesthesia, the pig was euthanized by controlled manual IV bolus overdose of Euthasol^®^ (pentobarbital sodium and phenytoin sodium solution), and death was confirmed by auscultation of cardiac arrest.

Data analysis was performed in Amide 1.0.4. DICOM data was loaded and ROI analysis was performed by placing a cylindrical ROI over the abdominal aorta and an area of the liver without major blood vessels. The same size ROI was used for all scan sessions. Raw MRI signal intensity data within regions of interest were extracted from MR images and exported into Excel (Microsoft) and percent signal enhancement was calculated as (signal-baseline)/baseline. The area under the curve (AUC) was calculated using the trapezoidal method to determine the percent clearance by organ.

## Results

All MRI sessions were successful, in that the pig remained anesthetized and stable during the MRI, high quality data was acquired, and the pig was recovered (except for the last scan where the pig was euthanized as planned). Figure 1 shows MRI before and immediately after injection of contrast agent, as well as at 13 minutes post contrast agent injection (maximal hepatic signal enhancement for Eovist), for the Eovist injection in scan session 1, the Multihance injection in scan session 4, and the Magnevist injection in scan session 3. In general, image quality was fair. State of the art motion compensation sequences such as radial VIBE were not available at our veterinary facility. Small motion artifacts are visible in the image as both blur and decreased signal to noise, and became more evident as the animal grew larger. High signal enhancement in the abdominal aorta following injection is clear in all cases. Hepatic signal enhancement is seen for Eovist and Multihance, with higher enhancement for Multihance than Eovist.

**Figure 1:**
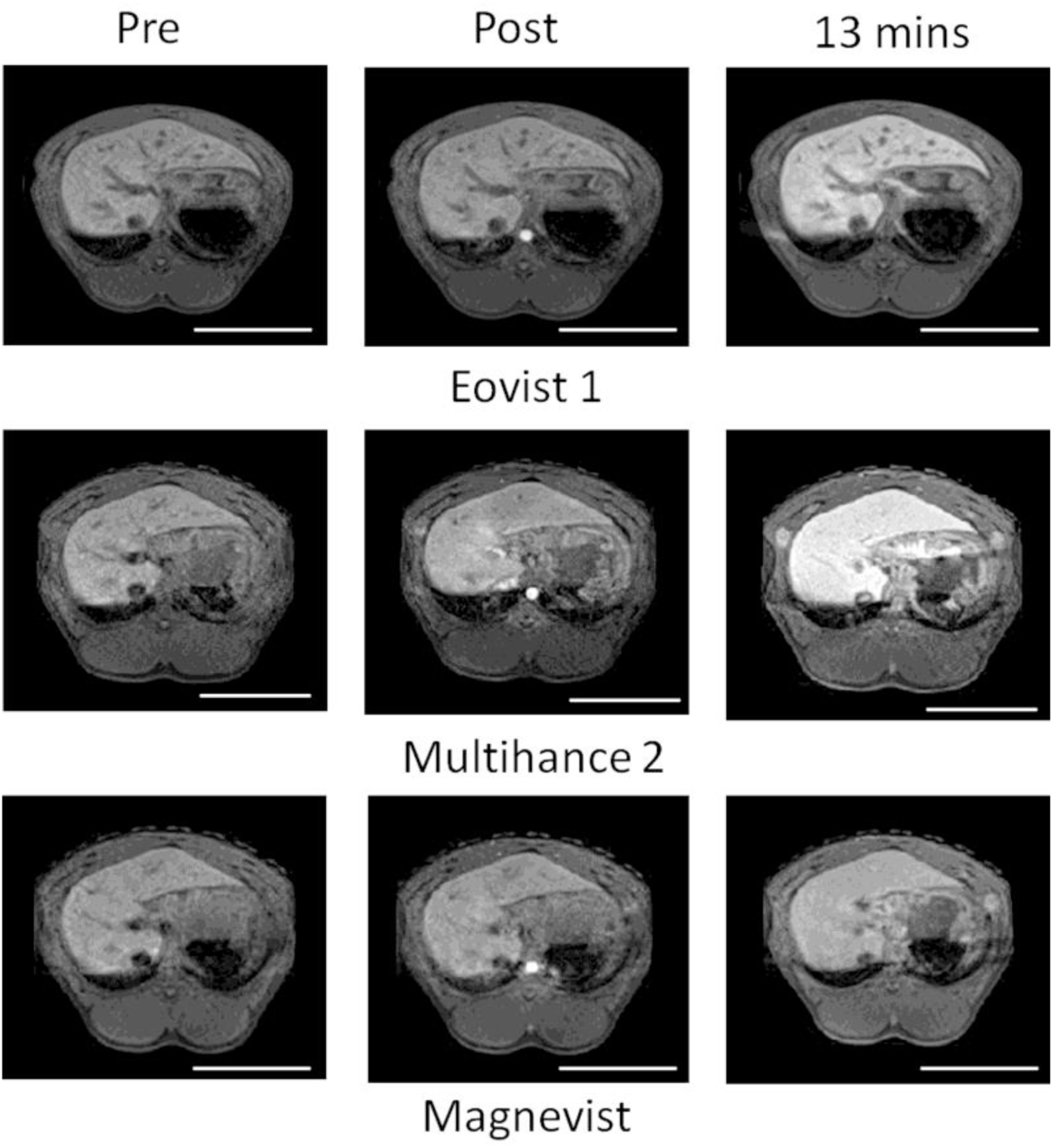
MRI of pig pre-, immediately post- and 13 minutes post-administration of Eovist (scan trial #1), Multihance (scan trial #2) and Magnevist. Scale bar is 10 cm.

Figure 2 quantifies the imaging data and plots normalized signal enhancement versus time in the abdominal aorta and in the liver. Table 2 lists the AUC for this data along with the CBT (°F) measured at the end of the experiment. For Eovist, both scans yielded nearly identical data, despite being performed first and last in the scan sequence and with a body weight increase of 30 kg over 5 weeks. Peak hepatic signal enhancement of 40% over baseline occurred at ∼12 minutes, with gradual signal decline to 20% enhancement at 60 minutes post injection. Liver AUC were nearly identical. Distribution of Eovist occurred rapidly in the blood, especially the second Eovist trial (scan session 5 in Table 1), on the same time scale as Magnevist. CBT for the two Eovist trials were similar.

**Table 2:** Summary of quantitative analysis of DCE curves as well as core body temperature at end of imaging session.

|  | Eovist 1 | Eovist 2 | Multihance 1 | Multihance 2 | Magnevist |
| --- | --- | --- | --- | --- | --- |
| Time at peak enhancement (seconds) | 666 | 680 | ND | 612 | ND |
| AUC Aorta | 1441 | 1546 | 3847 | 4900 | 2641 |
| AUC Liver | 1147 | 1136 | 1677 | 2150 | 514 |
| Core body temp (F) | 103.5 | 102.5 | 107.2 | 98.8 | 100.8 |

**Figure 2:**
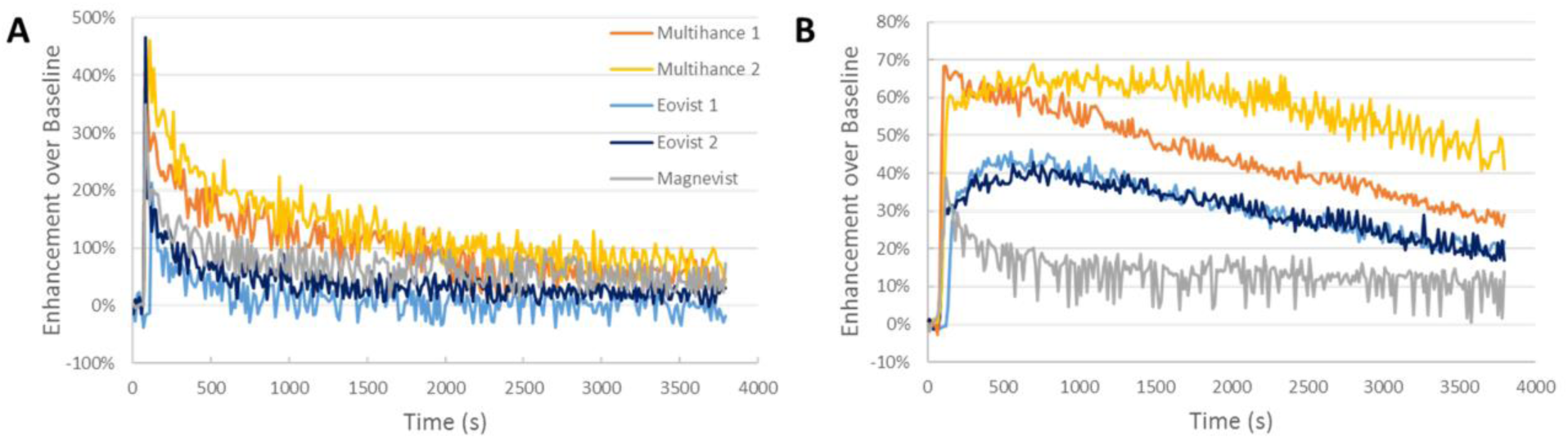
Plots of signal intensity enhancement over baseline versus time for A) abdominal aorta and B) liver parenchyma, for the five described experiments. Color key in A is same as for B.

For Multihance, there are two different results. In the first Multihance trial (Multihance 1, scan session 2 in Table 1), a very rapid hepatic signal enhancement is seen, with a rapid, linear decrease in signal intensity for the rest of the scan session. The CBT after the scan session was 107.2°F. In the second Multihance trial (Multihance 2, scan session 3 in Table 1) the addition of a cooling pad resulted in 98.8°F CBT and MRI results showed peak hepatic signal enhancement of 65% enhancement over baseline at ∼11 minutes post injection, with gradual signal decline to 45% enhancement at 60 minutes post injection. Distribution of Multihance in the blood occurred rapidly as with Eovist.

Clinical chemistry blood serum analysis showed that alkaline phosphatase (ALP) was slightly elevated during the entire experimental period, compared to normal values. Creatine kinase (CK) was highly elevated at scan session 2, 3 and 4 versus 1 and 5 but without control values for comparison. All other values were within normal ranges. (Supplemental Table 1).

## Discussion

In this study, we show for the first time that porcine liver has rapid accumulation of the two FDA-approved hepatospecific MRI contrast agents, Eovist and Multihance, following IV injection of equivalent human doses. For Eovist, the pig data is very similar to that seen in humans, both in terms of time to peak signal enhancement (∼10 minutes) and clearance rate from the liver to the bile (∼50% in 1 hour) ^18^. The porcine MRI enhancement following Multihance administration differs with humans, in that for humans, Multihance uptake by the liver is slow and inefficient, being mostly cleared by renal pathways. The pig results are similar to those previously observed in dogs, both for Eovist and Multihance, and are unsurprising given that both pigs and dogs express the same uptake transporter in hepatocytes, OATP1B4.

A major difference in scanning research animals versus companion animals is the type of information usually desired from the study. A clinical companion animal study employing hepatospecific MRI contrast agents likely seeks to identify tumors in the liver, or liver dysfunction. A research animal scan can of course accomplish this as well, but often, kinetic parameters of contrast agent influx and efflux, perfusion, or other biophysical parameters are desired, either in the context of disease or normal biology. So, while for both research and clinical animal scans, proper animal health must be maintained for the sake of the animal, there is a crucial need for proper animal health maintenance for reproducibility and accuracy of the data for research purposes. We observed that in this study, where for the two Eovist trials, despite being separated by five weeks and animal growth of 30 kg, keeping animal CBT within 1°F resulted in nearly identical temporal data of contrast agent uptake in the liver. This is contrasted with the two trials using Multihance, separated by only 1 week, where two very different temporal profiles were observed. We hypothesize that this was a result of overheating in Multihance 1 scan session - a 8.4°F CBT difference from Multihance 2 scan session. The temperature dependence of hepatospecific MRI contrast agent influx and efflux from the liver is well known in mice, with higher temperature resulting in faster uptake and clearance of agents from the liver ^25^. This is apparently the case in swine as well.

There are some drawbacks to this study and lessons learned. First, only a single pig was imaged, albeit 5 different times. Thus, it is impossible to make generalizations about uptake kinetics of contrast agents in all pigs as there are different strains of pigs, just as there are different strains of mice and rats. Second, a comparison between uptake kinetics of each agent cannot be made as the dose of contrast agent used in this study was different. The human clinical doses of Eovist (0.025 mmol/kg) is ¼ that of Multihance (0.1 mmol/kg). A second study with equivalent dosing is required to make comparisons between contrast agent uptake kinetics. Third, we used an older VIBE sequence with Cartesian k-space sampling, resulting in somewhat noisy images and motion artifacts, particularly as the animal aged. Newer motion compensated abdominal imaging sequences with radial k-space sampling should mitigate these issues ^26, 27^. Fourth, and perhaps not exactly a drawback, this study highlights a challenge in using domestic pigs for longitudinal studies. At purchase, the 2.5 month old pig weighed 25 kg and at the conclusion of the six-week study, the pig weighed 56.5 kg. This presents logistical challenges in performing serial imaging experiments on pigs past 5 or 6 months old that in some cases can be alleviated by using minipigs instead.

## Conclusion

DCE-MRI of swine following IV injection of hepatospecific MRI contrast agents Eovist and Multihance revealed rapid distribution of contrast agent in the blood, and high accumulation of contrast agent in the liver, shortly after injection. There was no retention of the non-specific agent Magnevist in the liver indicating it is not transported by hepatic OATPs, as expected. Proper animal health maintenance, especially temperature, seems essential for accurate and reproducible results. Given the disparity in contrast agent uptake kinetics with humans for Multihance, Eovist should be used in porcine models for biomedical imaging.

## SUPPORTING INFORMATION

**Table S1.**
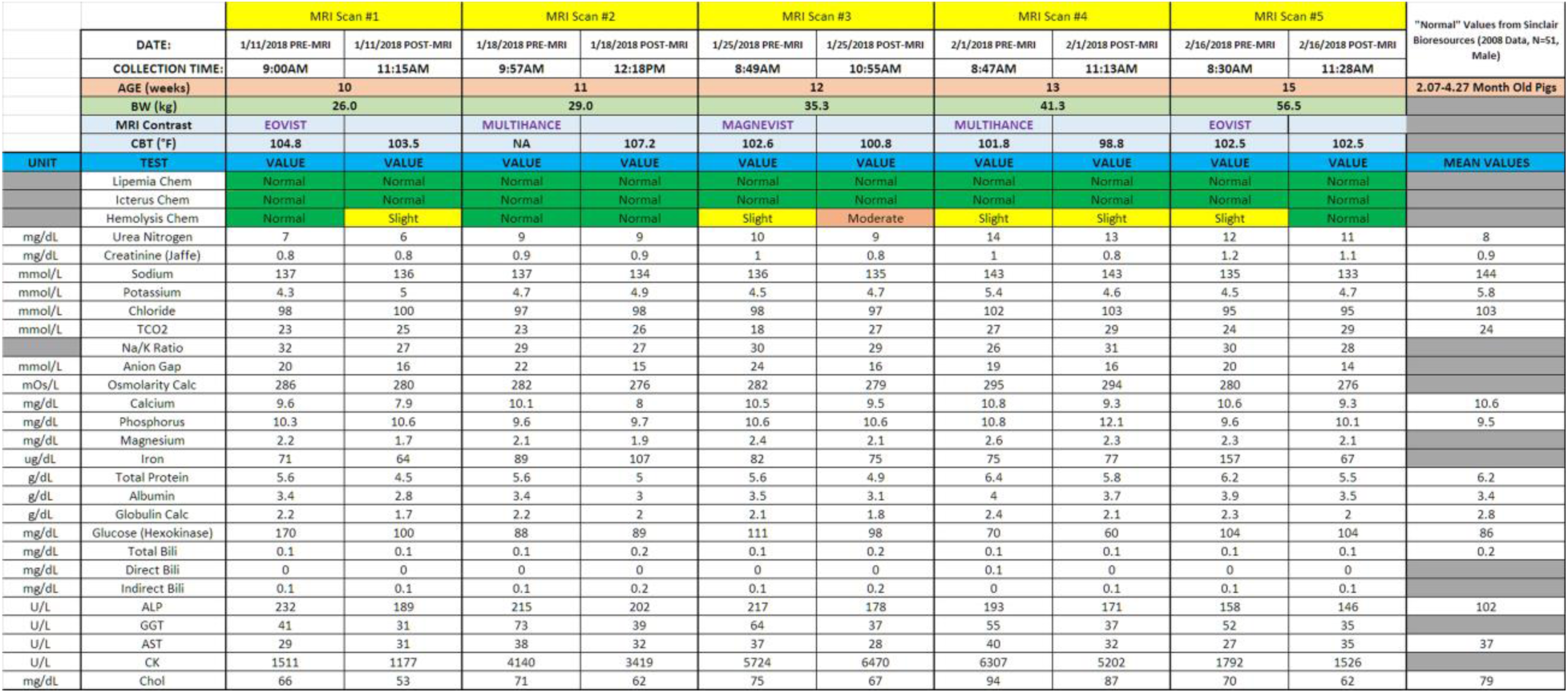
Blood chemistry and other animal information for all scan sessions.

## Acknowledgements

We are grateful to: MSU CVM Radiology technicians Rex Miller and Alexis Willis-Redfern for operating the CVM MRI, MSU RATTS team for animal procedures, MSU Campus Animal Resources for animal care and housing, Colleen Hammond for assistance with veterinary MRI protocols and Dr. Bruno Hagenbuch, University of Kansas Medical Center, for helpful conversations about OATPs. Grant support from the National Institutes of Health is acknowledged (NIH R01 DK107697 to EMS).

